# Neural Kernels for Recursive Support Vector Regression as a Model for Episodic Memory

**DOI:** 10.1101/2022.02.22.481458

**Authors:** Christian Leibold

## Abstract

Retrieval of episodic memories requires intrinsic reactivation of neuronal activity patterns. The content of the memories are thereby assumed to be stored in synaptic connections. This paper proposes a theory in which these are the synaptic connections that specifically convey the temporal order information contained in the sequences of a neuronal reservoir to the sensory-motor cortical areas that give rise to the subjective impression of retrieval of sensory motor events. The theory is based on a novel recursive version of support vector regression that allows for efficient continuous learning that is only limited by the representational capacity of the reservoir. The paper argues that hippocampal theta sequences are a potential neural substrate underlying this reservoir.The theory is consistent with confabulations and post-hoc alterations of existing memories.

## 1 Introduction

To retrieve episodic memories, brains need to elicit robust internal sequences of neuronal activity patterns that are linked to previous sensory-motor experiences. Thus, neural processes need to be in place that form such activity sequences as well as link them to sensory-motor areas while learning. Episodic memories are further known to be able to change over time by reconsolidation (Sara, 2000; Nader et al, 2000; Milekic and Alberini, 2002; Alberini and Ledoux, 2013), eventually even leading to false memories of events that never happened (Loftus, 1992; Hyman Jr. et al, 1995). This suggests that the architecture of episodic memory is versatile and local in time in the sense that any pair of memory items can be connected into a memory episode independent of context.

Since electrophysiological recordings in animals prohibit correlating activity sequences to introspective retrieval of episodic memories, memory-related activity sequences are typically studied in rodents in association with behavioral performance in navigational tasks (Lee and Wilson, 2002; Karlsson and Frank, 2009). Activity sequences of hippocampal place cells thereby have been reported to correlate to (Lee and Wilson, 2002; Dragoi and Buzski, 2006; Foster and Wilson, 2007) and to causally explain (Jadhav et al, 2012; Fernndez-Ruiz et al, 2019) memory-dependent navigation. Sequences have furthermore been found to exist even before a specific spatial experience has been made by an animal (Dragoi and Tonegawa, 2011, 2014; Farooq and Dragoi, 2019), suggesting that, at least part of the learning process is about establishing synaptic connections between existing intrinsic neuronal sequences and the sensory-motor areas that represent the content of the memory episode.

The idea that multi-purpose intrinsic neuronal dynamics is used to represent time series of extrinsic events has been invented multiple times under the names of echo-state networks (Jaeger and Haas, 2004), liquid computing (Maass et al, 2002) and reservoir computing (Jaeger, 2005; Schrauwen et al, 2007; Lukoeviius and Jaeger, 2009) and has proven to be both computationally powerful and versatile (Maass et al, 2002; Sussillo and Abbott, 2009), particularly, since multiple output functions can be learned on the same intrinsic activity trajectory (sequence) and played out in parallel.

There has been considerable previous work on how to construct a dynamical reservoir via the dynamics of neuronal networks (Haeusler and Maass, 2007; Sussillo and Abbott, 2009; Lazar et al, 2009). Also different learning rules for the synapses from the sequence reservoir to the output neurons were successfully explored, such as the perceptron rule (Maass et al, 2002), a Hebb rule (Leibold, 2020) or recursive least squares derived rules (Williams and Zipser, 1989; Stanley, 2001; Jaeger and Haas, 2004; Sussillo and Abbott, 2009). The more general applicability of reservoir computing to neuroscience is, however, still limited because several open questions remained, particularly about how to relate reservoir computing ideas to neurophysiological data: For example, can sufficiently rich reservoirs be realized with spiking neuronal networks? How can be found out whether reservoir spiking activity bears meaningful representations in the sense of Marr’s second level – as opposed to just being a “liquid” black box? Can a regression type learning rule be neuronally implemented using local Hebbian principles? How can new information (including false memories) be added to an existing episode or specific memory items be deleted? Can physiologically plausible models realize the universal approximation property Grigoryeva and Ortega (2018), or are the limits of learning already imposed by interference of weight updates at output synapses below the capacity limit (Amit and Fusi, 1994)? Particularly the latter problem is of fundamental importance in applying reservoir computing ideas to brain activity data, since available recordings (and also most models) are generally restricted to only a relatively small number of neurons (limiting representational capacity), whereas a whole real brain has close to infinite capacity for all practical (here experimental) purposes. Finding a representation of reservoir activity would thus eliminate capacity limitations and allow for efficient representation of a huge set of sensory-motor experiences in the synaptic weights.

Here, I propose a neuronal implementation of recursive kernel support vector regression as an *efficient one-shot* learning rule that is only limited by the representational capacity of the dynamical reservoir and allows for importance scaling (as compared to only graceful decay). Kernels thereby allow reservoir activity to be interpreted as representations in the sense of Marr (Hermans and Schrauwen, 2012), which adds to theoretical neuroscience by allowing for specific interpretations of neural activity. For example, I will argue that theta sequences of hippocampal place cells (Foster and Wilson, 2007) implement a kernel that represents distance in time or space and that the integration of auditory nerve activity at different delays implements a kernel representing time for acoustic stimuli in a cochlear frequency band. Below the capacity limit, the learning rule implements the well-known recursive (Gauss-Legendre) least mean squares (or FORCE) rule (Sussillo and Abbott, 2009) on the underlying neuronal patterns, showing that FORCE-learning is only limited by the capacity of the simulated or measured reservoir.

## 2 Results

Let us consider an episodic experience to be fully reflected by the sensory-motor evoked summed postsynaptic potentials 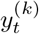 at all involved neurons *k* for all points in time *t*. In order to store the episodic experience as a memory, the synaptic inputs 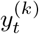 need to be linked to a pre-existing reservoir state ***x***_*t*_ such that, whenever ***x***_*t*_ is present afterwards, the learned synaptic connections ***w***^(*k*)^ from the reservoir evoke the same depolarizations

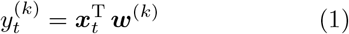

without the presence of the original sensory-motor activity (Fig. 1), i.e., ***w***^(*k*)^ solve a regression problem with ***x****t* as regressors. Considering only depolarizations *y*^(*k*)^ in such a one-layer feedforward network, one does not need to consider non-linearties during spike generation and, with reasonable approximation, the model is an effectively linear network. It also should be noted that in this paper I do not intend to explain the nature of the preexisting sequences ***x***_*t*_, and just assume that they exist. For the sake of simplicity, I further drop the neuron index *k*, since all considerations trivially generalize to multiple neurons.

**Fig. 1.**
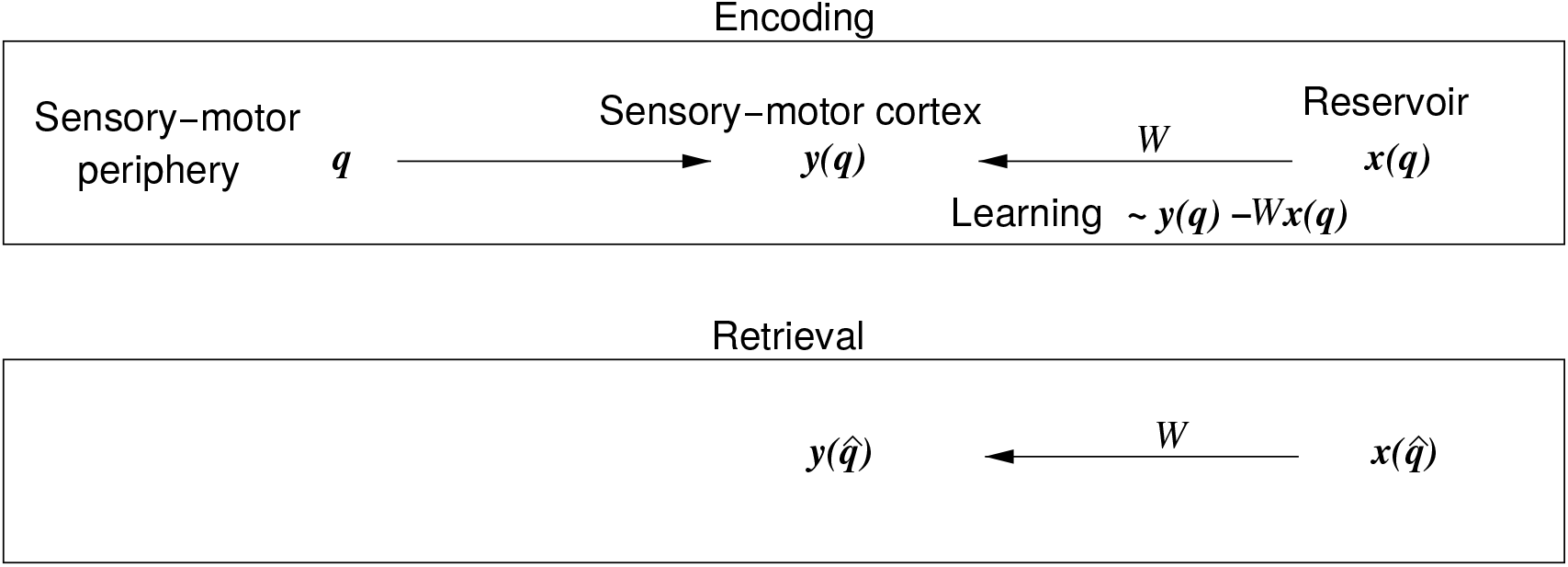
Conceptual overview. At any instance in time, let us consider ***q*** to encode any sensory-motor experience of an agent (human, animal or machine). A neocortical representation ***y***(***q***) of this experience is evoked by the sensory afferents, and motor efference copies. The current state ***x*** of a reservoir (e.g. in the hippocampal formation) is linked to the temporally coincident experience ***q*** by synaptic learning of the connections *W* from the reservoir to the neocortex. The synaptic change is thereby proportional to the error signal ***y*** − *W* ***x***; see eq. (5). During retrieval, the reservoir state ***x*** previously associated with a real, or confabulated experience 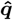, evokes a corresponding neocortical representation ***y***(***q***).

Besides the scalar product in eq. (1), biological feasibility imposes two more constraints on how one models learning. First, synaptic plasticity should be activity-dependent and therefore the weights should be a superposition of existing neuronal activity patterns,

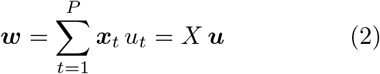

with *X* = (***x***_1_, …, ***x***_*P*_) (see Discussion on representer theorem). Second, the learning rule needs to be recursive, i.e., new input-output pairs (***x***_*P* +1_, *y*_*P* +1_) should be added such that eq. (1) holds for all previous patterns (no interference) until the capacity limit and memory decay beyond the capacity limit should be importance based. In short, the learning rule is supposed to identify the loads ***u*** such that the outputs *y*_*t*_ are exactly recovered by the model,

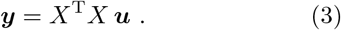

As long as the kernel matrix *K* = *X*^T^ *X* is invertible (below the capacity limit), the solution for ***u*** is exact and straight forward. For noninvertible or badly conditioned *K* (at or above the capacity limit), the standard approach would be to use the pseudo inverse *K*^∗^ of *K*, which optimizes the mean squared deviation between output ***y*** and model output *K K*^∗^ ***y*** and leads to the classical recursive least squares (RLS) algorithm if applied recursively. RLS on the loads ***u***, however, has two main disadvantages. First, RLS makes explicit use of time making it hard to modify memories by post-hoc insertion of new detail within an existing memory sequence. Second, RLS on the loads ***u*** is hard to interpret biologically.

I therefore suggest, as an alternative approach, to solve the regression problem by maximizing

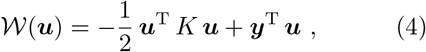

which, for invertible *K*, yields the exact recovery condition eq. (3), therefore justifying the use of 𝒲 as the underlying objective function. Moreover, the maximization problem from eq. (4) can be derived as the dual problem of support vector regression for *ε*-insensitive loss (see Methods and Vapnik, 1995; Schölkopf and Smola, 2002), further supporting the interpretation of regression.

Since support vector approaches translate to non-linear models using the kernel trick 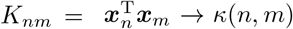 (Vapnik, 1995), the model also provides a foundation for neural implementations of kernels, which can be considerd as *representations* of the topological space spanned by *n* and *m*. In the same sense as Marr saw representations to be connected to the algorithmic level, the kernel represents the space of *n* and *m* in a sufficient way to fully specify the outlined regression algorithm and thus, following (Hermans and Schrauwen, 2012), I suggest to consider it being true neural representation of this in contrast to considering representations as activity patterns in undersampled cell populations.

Maximizing 𝒲 results in an update rule for ***u*** (see Methods) that translates into a weight change **Δ*w*** = ***X*Δ*u*** of

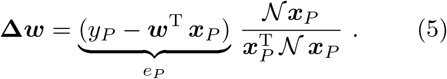

with 𝒩 = 𝟙 − *X K*^−1^ *X*^T^, and an iteration rule

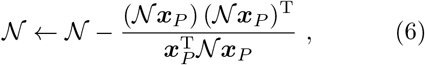

that is equivalent to RLS without forgetting (i.e., without regularization). The learning rule is one shot in the sense that, for any new pattern, the update rules have to be applied only once and it allows for the functional interpretation *error* (*e*_*P*_) times *novelty* (*𝒩*): Because 𝟙 − 𝒩 is a projection operator (see Methods), *𝒩* ***x***_*P*_ will be 0 whenever ***x***_*P*_ equals one of the previous patterns already included in *X*, whereas any component of ***x***_*P*_ that is orthogonal to all patterns in *X* will be unaffected by 𝒩. The action of 𝒩 can thus be computationally interpreted as *novelty detection*. For a naive learner (*P* = 0), the rule is plain Hebbian, since the error equals the output and the novelty equals the input pattern. In the Discussion, I will suggest a biologically feasible implementation of 𝒩 and its learning as anti-Hebbian updates of a recurrent neural network. Importantly, the translation into neuron space resulting in eqs. (5,6) is only required to show how the learning rule can be biologically implemented. In contrast to RLS, it is not necessary to use these update rules for all ensuing applications, which are only relying on the numerically much more tractable update rule for the loads ***u*** presented in eq. (7) in the Methods section.

As a first neuroscience application, I refer to hippocampal theta sequences (Fig 2A): Roughly, one considers a subset of place cells to fire in sequence in every cycle of the hippocampal theta oscillation of the local field potential (about 8 Hz in rodents). In the subsequent cycle, the starting neuron of the previous cycle drops out of the sequence but a new neuron is added at the end of the sequence. Thus the activity patterns of close-by cycles are similar, whereas they become more and more distinct the further the cycles are spaced apart. In the simple theta sequence model outlined above, the overlap (scalar product) of activity patterns decays linearly (see Methods) implementing a kernel *K*_*mn*_ = *κ*(*n* − *m*) as a function of the distance *n* − *m* of the two cycles (Fig. 2B).

**Fig. 2.**
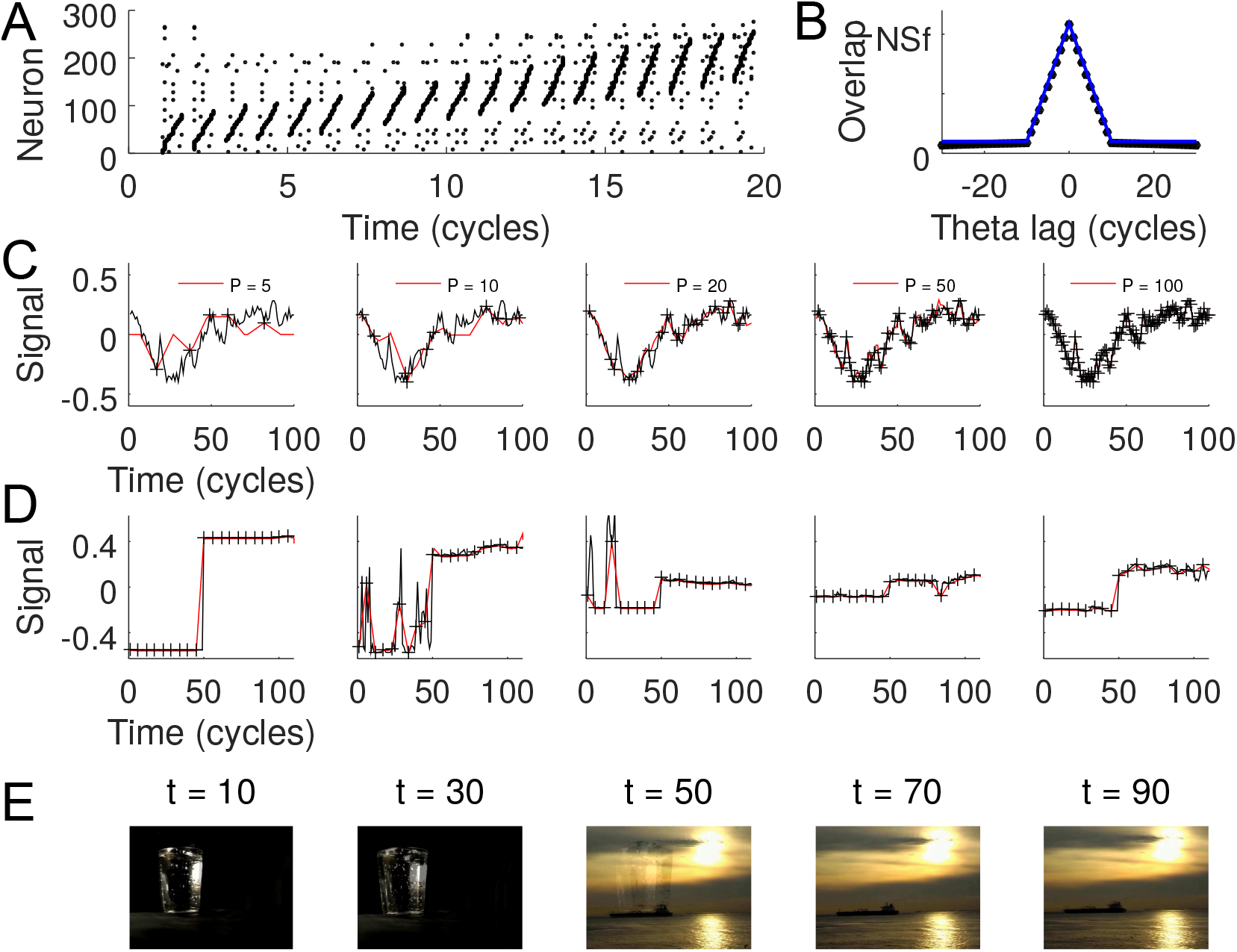
Episodic learning with theta sequences. (A) Spike raster plot of the first 300 of *N* = 10, 000 neurons implementing theta sequences as described in Methods section *Theta sequences* (sparseness *f* = 0.01, sequence length *S* = 10). In every theta cycle the sequence moves one neuron upward. (B) Kernel derived as scalar product between population patterns from the simulations shown in A (black dots) and theoretical prediction (blue line). (C) Retrieval (red line) of a low-pass noise signal (black) of length *T* = 100 from *P* observations (crosses; for *P* see insets from left to right) using the theoretical kernel from B. The signal was generated as a running average (50 time steps) of white noise. (D) Same as B. Brightness signal for five example RGB channels from a movie scene (*P* = 20, *T* = 111, *N* = 576 *×* 768 *×* 3). (E) Retrieval of movie snippet (five example frames shown) from .*re_potemkin*, a copyleft crowd sourcing free/open source cinema project (https:/re-potemkin.httpdot.net/). Original movie snippet and reconstruction are provided as Videos S1 and S2.

Inserting the triangular linear kernel from Fig. 2B into the learning rule derived by recursively maximizing *𝒲*, one can recover the original signal *y*_*t*_ without simulating the underlying reservoir. Increasing detail of the original signal can be retrieved the more pairs (***x***_*t*_, *y*_*t*_) one takes into account for learning (Fig. 2C). Since the kernel is a continuous function, the capacity has become infinite, i.e., any function *y*_*t*_ can be recovered if the neuron number *N* becomes infinite.

As mentioned above, generalization to multiple neurons is trivial, and to illustrate let us consider each output neuron to reflect one RGB color channel of any pixel in a movie (1.3 million neurons). Using only 20 of 110 movie frames already allows for recovery of the movie snippet with a compression below 20% (Figure 2D,E).

By construction, the learning rule has no explicit dependence on time, thus the order in which pairs (***x***_*t*_, *y*_*t*_) are presented makes no difference to the final fit (Figure 3A), which is not the case for the FORCE rule derived from classical least squares. Biologically, this means that any episode can be posthoc modified by learning new pairs (***x***_*t*_, *y*_*t*_) with temporal contingencies reflected in the kernel arguments, generating a model of false memories (Figure 3B).

**Fig. 3.**
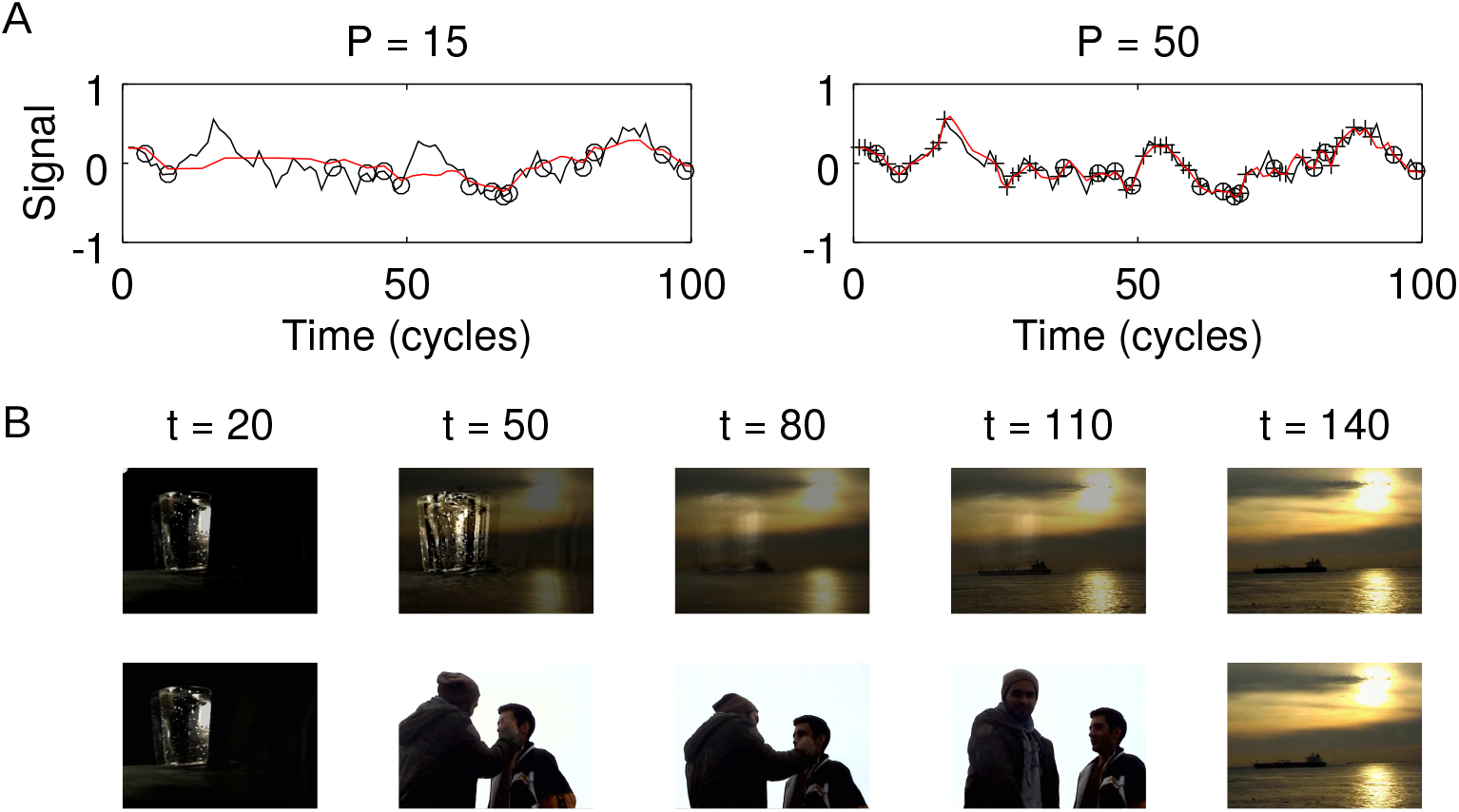
Post-hoc addition of memory items. (A) Left: Retrieval (red) of a low-pass noise signal (see Fig. 2C) of length *T* = 100 for *P* = 15 randomly positioned inputs (circles). Right: Same as left after 35 further inputs (crosses) have been iteratively added to the learning process. (B) Illustration of A for posthoc insertion of a movie scene. Top: original movie sequence (*P* = 20). Bottom: Movie sequence after a new scene has been inserted to the original snippet (*P* = 35). Movies are provided in Videos S3 and S4.

Every memory system is finite and the way of forgetting fundmentally determines its usefulness for practical applications. A graceful decay of memories over time (Amit and Fusi, 1994) is already quite an advantage to catastrophic forgetting in attractor networks (Hopfield, 1982), however, the behavioral relevance of a memory may not just depend on how old or young it is. I therefore introduce an importance scaling into the learning rule in that loads *u*_*t*_ are multiplied with some attentuation factor 0 *≤ a*_*t*_ *≤* 1. Thus, if one chooses *a*_*t*_ = *λ*^(*T*−*t*)^, 0 *< λ <* 1 one retains a graceful decay over time as in standard RLS. The resulting learning rule that maximizes modified 𝒲 is then obtained by only the small modification of replacing the kernel *κ*(*n, m*) by *κ*(*n, m*) *a*_*n*_ *a*_*m*_ (see Methods). The effect of importance scaling is illustrated in Figure 4A,B, where the learning rule is told to pay more attention to a certain time interval at the cost of worse reconstruction in other time intervals.

**Fig. 4.**
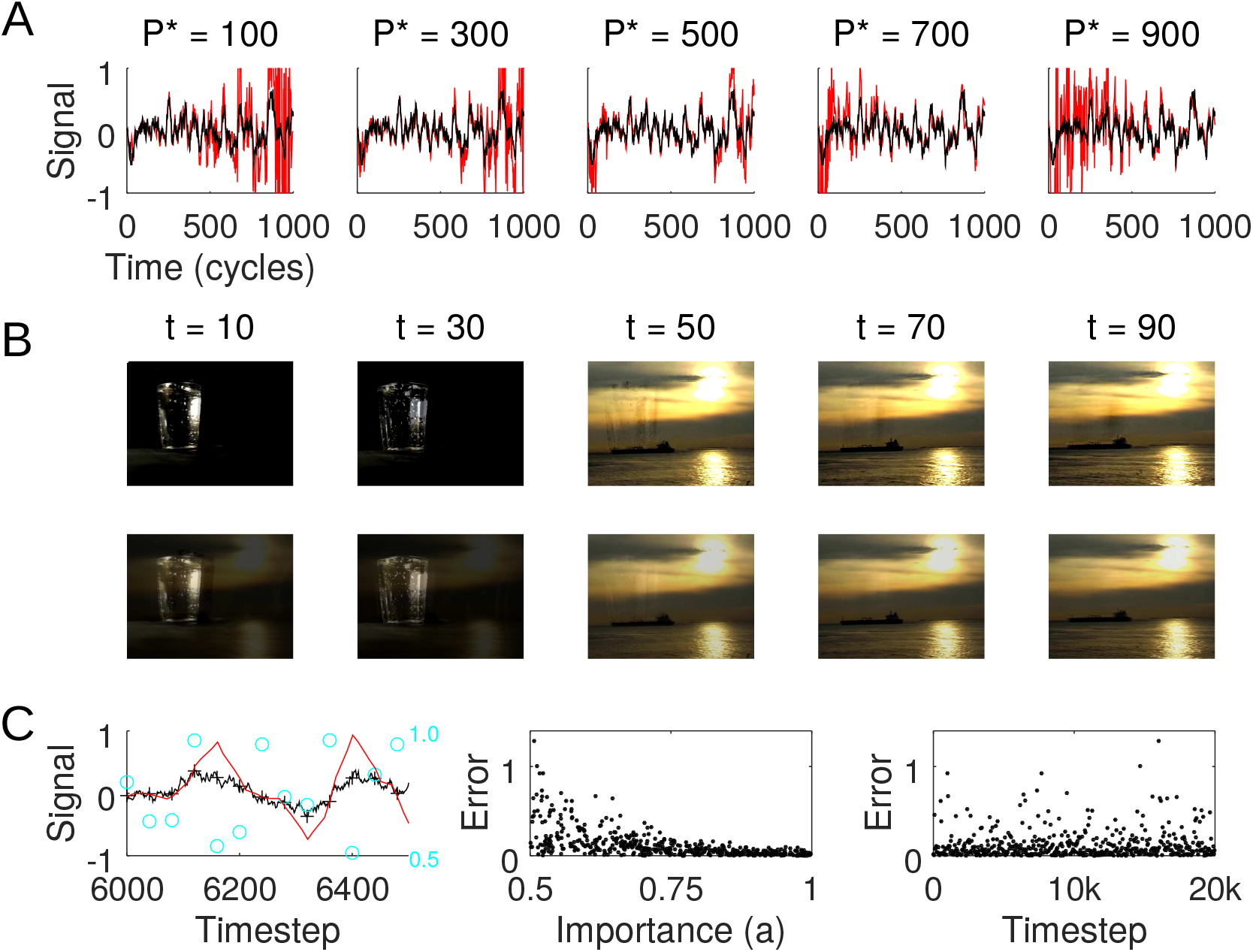
Importance scaling. (A) Retrieval (red) of a low-pass noise signal (black; see Fig. 2C) for attenuation parameters *a*_*t*_ = *λ*^|*P*∗−*t*|^ with *λ* = 0.999 and varying importance centers *P* ^*∗*^ (see titles). (B) Illustration using the movie snippet from Figure 2 with importance in the beginning (*a*_*t*_ = *λ*^*t*^,top) and in the end (*a*_*t*_ = *λ*^110−*t*^, bottom). In the image sequence on top one stills sees a erroneous reflection of the glass in the last three images, whereas in the bottom sequence the glass in the first to frames shown erroneously displays the yellowish colors from the end. Movies are provided in Videos S5 and S6. (C) Left: Retrieval (red) of a low-pass noise signal (black) with *N* = 20, 000 time steps (only shown between time step 6,000 and 6,500) and *P* = 500 patterns (crosses) with random importance values *a* (cyan) between 0.5 and 1. Middle: Reconstruction error (absolute difference between black an red line) negatively correlates with *a* for all *P* = 500 patterns. Right: Error has no dependence on time.

Importance may randomly vary over time and thus temporal contingency in *a* values should not be a necessary prerequisite for importance scaling. Applying the learning rule in a scenario with random *a* values shows that retrieval error is indeed largest for small *a* independent of time (Figure 4C). Posthoc increase of *a* could thus be considered as a model of memory consolidation, posthoc decrease of *a* as a model of extiction learning.

With importance scaling as a weighting mechanism at hand, let us now revisit the original capacity question. In the language of the recursive updating rules from equations (7) and (8) the memory and computational demand scale with square of the number of patterns *P*. A straightforward choice to limit the capacity is to introduce a cut-off dimension *d*_*c*_ such that only the *d*_*c*_ patterns with highest importance values *a* are stored in the algorithm and the other dimensions are set to 0. In Figure 5A,B we vary *d*_*c*_ for low-pass filtered noise signals of different length with linearly increasing importance towards the signal end and observe that for low *d*_*c*_, the reconstruction error increases relatively soon, whereas for *d*_*c*_ ⪆ 300 reconstruction worked well even for signal lengths up to 10 times larger then *d*_*c*_, which reflects that the geometry of the kernel fits the correlational structure of the signal.

**Fig. 5.**
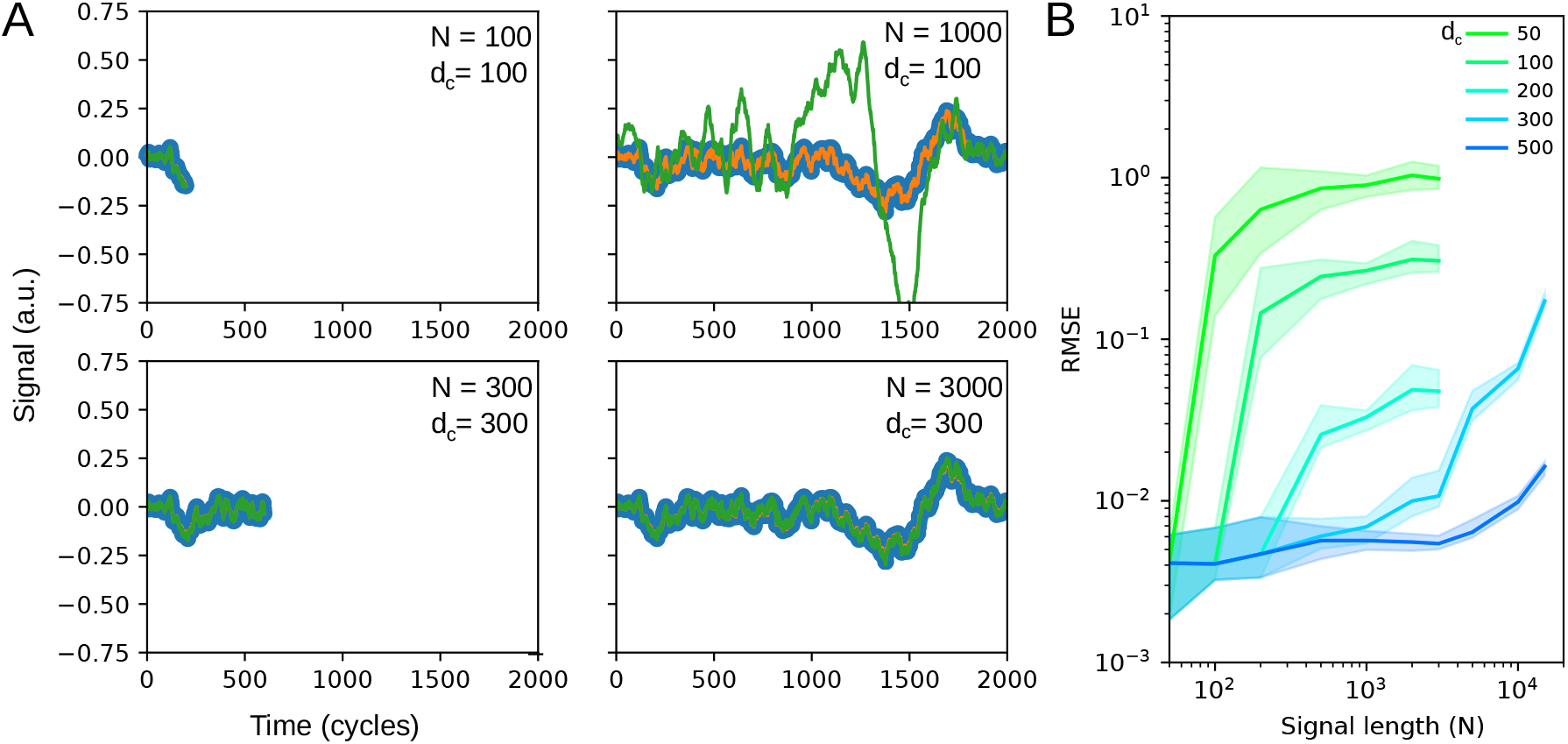
Capacity. (A) Example reconstructions (green) of a signal (orange) for smaller (top) and larger (bottom) cut-off dimensions (*d*_*c*_ = 100 and 300, respectively). Sample points used for reconstruction (signal length *N* = cycles / 2) are shown in blue. For the left panels the signal length equals the cut-off dimension. On the right the signal is 10 times longer then the cut-off (only 1000 data points shown). (B) Reconstruction error (root mean squared) for different cut-off dimensions (colors as indicated) as a function of signal length (solid lines indicate mean from 20 repetitions, shaded areas the 90 percent quantile). Results are derived from a low-pass noise signal with a running average over 100 time steps and a triangular kernel with length 25 time steps.

The need to adjust the kernel length to the time scale of signal fluctuations suggest that more specific signal properties require more specifically desigend kernels. In most neuroscience applications, sensory signals are not random but reflect physical constraints of the environment or the sensory periphery. As a next example I therefore consider functions with bandpass characteristics similar to cochlear frequency channels. Knowledge about the preferred local structure of a function (oscillations with a certain frequency) suggests a kernel with similar bandpass characteristics (see Methods and Figure 6A). In contrast to the triangular linear kernel which only represents temporal distance, the band kernels represent temporal distance (by their decay) and frequency. Learning is then performed on each cochlear frequency channel separately and the fitting benefits from both recovering the function values at a few points and the fine structure of the kernel. A posthoc synthesis across frequency channels recovers the original soundwave with high fidelity and smaller memory demand as the original sampling (see Methods section *Frequency kernels*).

**Fig. 6.**
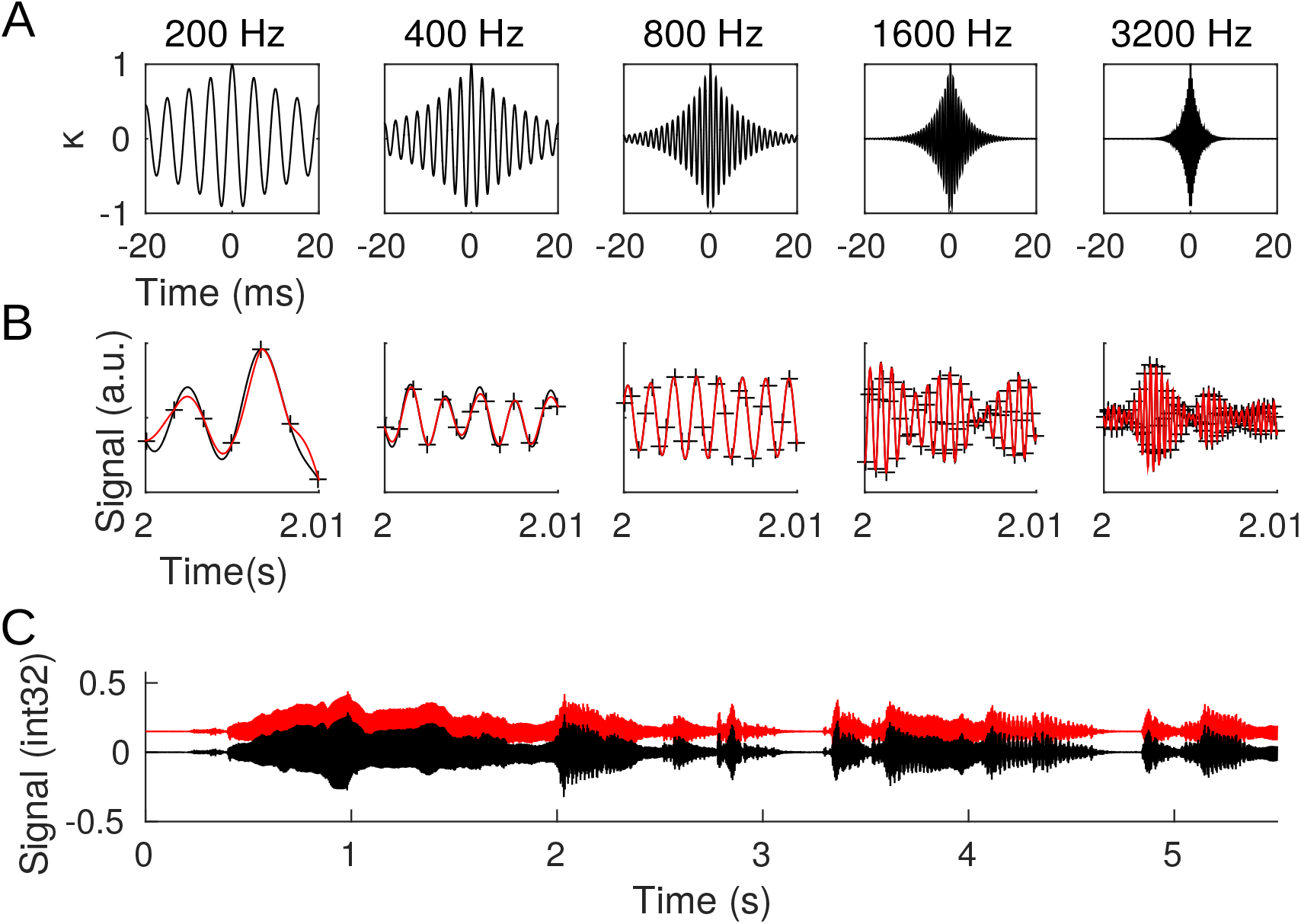
Sound reconstruction. (A) Kernels representing time in a frequency channel with center frequency on top (see Methods section *Frequency kernels*) (B) Retrieval (red) of the signal (black) in five of the frequency channels (crosses mark memory items). (C) Reconstruction (red; moved upward for reasons of illustration) of the original sound signal (black; the beginning of the song http://ccmixter.org/files/texasradiofish/63300, CC BY NC) by summing over the filter-weighted channel components (see Methods section *Frequency kernels*). Reconstructed sound file is provided in Audiofile S8, well as the identically filtered original sound wave (Audiofile S7)

## 3 Methods

### Recursive Support Vector Regression

Linear support vector regression with *ε*-insensitive loss (Schölkopf and Smola, 2002; Vapnik, 1995) is derived from minimizing the squared L_2_-norm 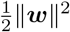 of the weight vector of the linear model *f* (*x*) = ***w***^T^ ***x*** + *b* under the inequality constraints 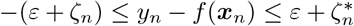, with 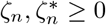, and including the sum of slack variables 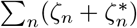 as a regularizer.

The classical work has shown that the resulting optimal solution yields a weight vector of shape

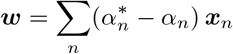

that maximizes the dual problem

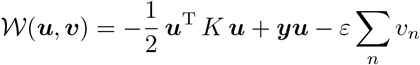

with 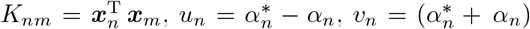 under the constraints *α, α*^∗^ *≥* 0. Hence, for every local maximum of *𝒲* regarding ***u***, there is a combination of 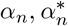 that minimizes ∑_*n*_ *v*_*n*_, i.e., *α*_*n*_ = 0 if *u*_*n*_ *>* 0 and 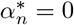 if *u*_*n*_ *<* 0. For *ε →* 0, the maximum in (***u, v***) converges to *α*_*n*_ = 0 or 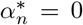, and thus, in this limit, one can drop ***v*** from the equations.

Here, a recursive learning rule is derived such that *𝒲* remains at this maximum if a new obervation (*y*_*p*_, ***x***_*P*_) is added. One therefore denotes 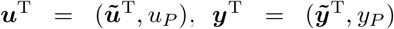, and 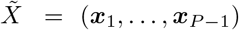 and find the optimum of

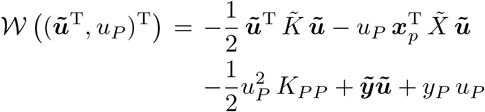

by

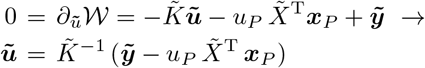

and

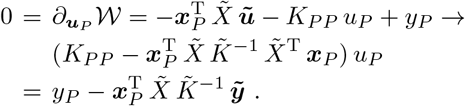

If one denotes the optimum loads of the previous *P* − 1 inputs by 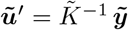, one can express the optimality conditions using 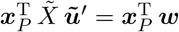, as

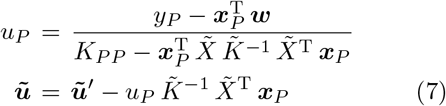

The update rules for ***u*** from eqs. (7) require computation of the inverse of 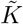, which, a) is computationally costly and, b) biologically not straight forward. I therefore derived an iteration rule using the Sherman-Morrison-Woodbury indentity Nocedal and Wright (2006), which yields an iteration equation for *K*^−1^ from the *P −* 1st to the *P* th pattern

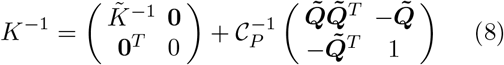

with 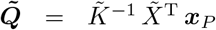 and 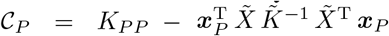. The iteration equation (8) can be proven by elementary algebra (*K*^−1^ *K* = 𝟙).

### Remarks

- Translation of update rules from eqs. (7) to weight updates **Δ*w*** = *X***Δ*u*** is straight forward:

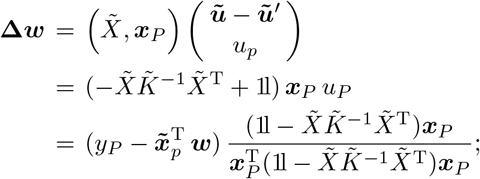

see Result from eq. (5).
- 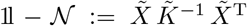, and 𝒩 are projection operators, since [𝟙 − 𝒩]^2^ = [𝟙 − 𝒩] and 𝒩^2^ = 𝒩.
- If *P −* 1 ≤ *N* and patterns are linearly independent, 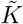 is a Gramian and, hence, invertible.
- For *P −* 1 exceeding *N*, 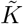 can no longer be exactly inverted. Formally this is not necessary using a kernel representation, since the kernel operates on an infinite dimensional Hilbert space. Biologically, for a finite number *N* of neurons, approximate inversion can be obtained by importance scaling (see below).
- Recursively adding data points continously increases the dimensions of the matrix *K*^−1^ and, hence, memory and computational costs. A brute force strategy to avoid this numerical divergence is to introduce a cut-off dimension, after which one removes the patterns with lowest importance values *a*. For all figures except Fig. 5, in which we explicitly study this parameter, we used a cut-off dimension of 300.

### Importance scaling

Importance is introduced by attenuation factors 0 *≤ a*_*t*_ *≤* 1 that scale the inequality constraints of Support Vector Regression: 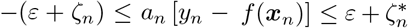. If *a*_*n*_ is small, slack variables can also be small and the the pair (*y*_*n*_, ***x***_*n*_) contributes little to the loss via the regularizer. The resulting optimal solution is very similar to the one without attenuation factors, only the weight vector are now

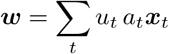

which, in the computation of the recursive learning rule, requires to replace

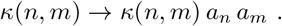

Biologically, this rule maps to an attenuation of the inputs ***x***_*t*_ *→ a*_*t*_ ***x***_*t*_. Thus, patterns with low *a*_*t*_ are treated as more different to patterns with large *a*_*t*_, even if they have similar structure.

The scaling of the kernel also has interesting consequences for situations in which the *K* is no longer invertible (*P > N*) if constructed from a finite population of neurons. In this case, one nevetherless, can apply the iteration equation (8), however, patterns with small *a*_*n*_ will contribute only little to 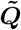 as the respective rows are scaled down in 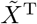. The resulting matrix is hence no longer an exact inverse, but the patterns for which the “inversion” fails mostly are those with low *a*_*n*_. This is best illustrated by assuming *a*_*n*_ = 0, in which case the pattern ***x*** has no contribution to 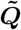 and hence *K*^−1^, as if it would not have been used for learning. Functionally modulating plasticity with *a* also allows a post-hoc improvement of an existing episodic memory, by setting higher importance *a*_*n*_ to this pattern if the epsiode is presented as second time.

### Theta sequences

Sparse binary random patterns ***ξ***_*n*_ with 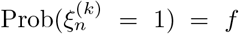 are assumed to represent hippocampal ensembles that fire together at a specific phase of the theta cycle. Given that *S* of those ensembles are activated in sequence during a theta cycle the population pattern in cycle *t* equals

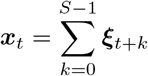

For a population of *N* neurons, the overlap of two such patterns can be computed as

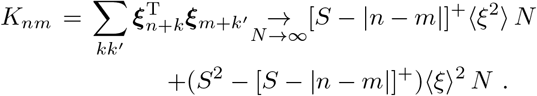

For independent binary random variables, one finds ⟨ *ξ* ⟩ =⟨ *ξ*^2^⟩ = *f*, and thus the overlap is a linear trangular kernel

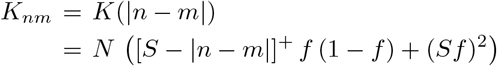

as depicted in Figure 2.

### Frequency kernels

The cochlea separates a sound *s*(*t*) into frequency channels that roughly act as band-pass filters and can thus be characterized by a filter kernel *γ*_*f*_ (*t*), with *f* denoting the center frequency of the cochlear channel. If one assumes multiple (*k* = 1, …, *N*) auditory nerve fibers to connect to such a frequency channel the linear response of each of those fibers can be modelled as 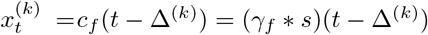 with a fiber specific delay Δ^(*k*)^ that may reflect differences in fiber lengths, diameters or myelination.

For a large number *N* of fibers the resulting kernel can be computed as an integral

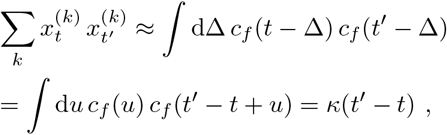

Which corresponds to the autocorrelation of cochlear response and, for long broadband signals *s* equals the autocorrelations of the filtes *γ*_*f*_. The exponentially decaying kernel used in Figure 6 reflects exactly such a prototypical autocorrelation.

Specifically, a sound signal (the beginning of the CC BY NC song *I’ll be your everything* by Texas Radio Fish, http://ccmixter.org/files/texasradiofish/63300) was passed through a gamma tone filterbank consisting of seven channels (center frequencies 2^*k*^ *×* 200 Hz, *k* = 0, …, 6) with width constants 2.019*ERB* (Glasberg and Moore, 1990). In each of the channels *ρ*_*k*_ data points per cycle (equally spaced) were selected for learning. The parameters *ρ*_*k*_ where channel (*k*-)dependent and equaled 6, 4, 3, 3, 1.5, 1, .25 for *k* = 0, …, 6. The recursive KSVR was fitted in each channel indepedently in chunks of 500 data points.

For full audio reconstruction, the reconstructed signals were Fourier-transformed in each band and divided by the Fourier transforms of the respective gammatone filter kernel omitting frequencies below 10 Hz and above 20 kHz. These filter-corrected components were backtransformed, summed and rescaled to the root-mean-square level of the original signal.

## 4 Discussion

Kernel support vector regression (KSVR) is a powerful tool for function fitting. Here, I presented a biologically plausible neural implementation of recursive KSVR that enables storing episodic memories as temporal sequences of retrieved sensory-motor activity patterns *y*_*t*_ (i.e., fitting *y*_*t*_). The kernels can be biologically interpeted as scalar products of activity patterns ***x***_*t*_ of a reservoir, and provide a neural representation of temporal distance.

Hippocampal theta sequences provide a well-known example that realize exactly such a reservoir. However, already in the hippocampus, neuronal activity not only consists of sequence-type activity, but also exhibits rate modulations induced by changes in the sensory environment generally known as remapping (Muller and Kubie, 1987; Leutgeb et al, 2005; Fetterhoff et al, 2021). Thus, behavior-related neuronal activity may always contain both, reflections *W* ***x***_*t*_ of the reservoir, and feed-forward sensory motor drive, thereby balancing expectations (i.e., reservoir-driven activity) and sensory reality. This combination of top-down and bottom-up input streams is widely considered to be a general design principle of the neocortex (Douglas and Martin, 2004; Larkum, 2013), resulting in sensory-motor activity patterns *y*_*t*_ at the same time reflecting stimulus-driven responses and intrinsic dynamics as, e.g. reflected by synfire chains (Abeles et al, 1993).

While the view of neocortex as a hierarchical combination of sensory-motor prediction loops (Ahissar and Kleinfeld, 2003) is probably a good proxy of neurobiological substrate, it is not widely explored in classical artifical neural network research. There, the universal approximation theorem, as a hallmark result, states that neural networks can approximate any function to arbitrary degree of precision (Cybenko, 1989; Hornik, 1991) which rather views brains as feed-forward function fitting devices. The field of reservoir computing has extended this idea towards the temporal domain by suggesting intrinsic neural dynamics to represent a time axis as the independent variable of function fitting (Jaeger, 2005) and thereby allows neural networks to generate predictions varying with time. However, to be able to operate on a continuous stream of sensory inputs, the learning rules for the output synapses of the reservoir need to be able to recursively update (Williams and Zipser, 1989; Stanley, 2001; Sussillo and Abbott, 2009), which requires a biological interpretation of the common least-mean square derived ideas.

Here, I suggest that the iterative update of the projection operation 𝒩 that only requires anti-Hebbian type outer products, can be implemented as anti-Hebbian learning of a simple recurrent neuronal network: In the neural space of synaptic weights, *X K*^−1^ *X*^T^ = 𝟙 − 𝒩 is of outer product form as seen from eq. (8). The matrix 𝒩= 𝟙 − *X K*^−1^ *X*^T^ can thus be interpreted as the connectivity of a recurrent neural network that is learned by anti-Hebbian updates, i.e.,

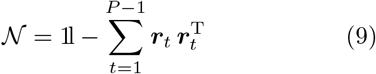

with 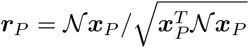. Since *X K*^−1^ *X*^T^ is a projection matrix (see Methods), one furthermore can write 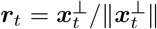 with 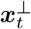 being the component of ***x***_*t*_ that is orthogonal to all previously learned patterns.

This leads to the following interpretation of ***r*** as the activity of a neural network in discrete time *s*

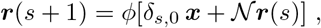

where the network is initialized at ***r***(*s* = 0) = 0, the input ***x*** is present only at time step *s* = 0, and *ϕ*(***z***) = ***z****/‖* ***z ‖***. As a result of this dynamics 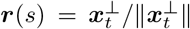 for all time steps *s >* 1. This dynamical fixed point state will then produce an anti-Hebbian weight update from eq. (9).

A further drawback of RLS-derived rules was their lacking theoretical foundation since they made explicit use of the reservoir patterns that, for technical reasons, were limited to a small sub-sample of neurons. Here, I use the generalized representer theorem (Schölkopf et al, 2001) to translate the weight update into an update rule for the loads (coefficients) ***u*** of the input patterns *X* and thereby avoid an explicit representation of the neural feature space ***x***_*t*_ and instead only require a kernel representation (Hermans and Schrauwen, 2012). Formulation of the learning rule on the loads allows analytical insights for reservoirs of size *N → ∞*, but also reduces the computational demand of simulating (or recording) from a large number of neurons.

Importantly, this paper considers reservoir activity only in the context of memory retrieval but not replay of reservoir sequences. Replay in the context of reservoirs has often been used to improve performance and stability (Mayer and Browne, 2004; Jaeger, 2010; Sussillo and Abbott, 2012; Reinhart and Jakob Steil, 2012; Laje and Buonomano, 2013; Jaeger, 2017; Leibold, 2020). However, changing reservoir patterns would require to also change the readout-matrix to maintain the originally learned memory traces *y*_*t*_ (Sussillo and Abbott, 2012; Reinhart and Jakob Steil, 2012; Jaeger, 2017). In the context of the model presented here, relearning is not necessary as long as the kernel remains fixed, i.e., the topology of the space is constant. Neurobiologically, however, such a trick would require to change the weights ***w*** by replacing the matrix *X*.

I presented two neurobiological examples of how kernel representations are or may be implemented, hippocampal theta sequences and auditory nerve fiber populations. Temporal sequences of activation patterns, however, are ubiquitous in sensory-motor systems and occur on multiple time scales. Thus the proposed theory may also apply to a multitude of other examples. A prerequisite is to find a continuous representation of time in the population patterns that then translates via a scalar product into kernels with continuous time depedence. Further such examples could be the long-term changes of the hippocampal rate code of place cells (Mankin et al, 2012; Ziv et al, 2013), activation of cerebellar purkinje cells during limb movements (Hewitt et al, 2011), or olfactory driven activity that evolves along fixed trajectories after odor presentation (Stopfer and Laurent, 1999).

## Supporting information

Wave files for Audio S7 & S8

## Supplementary information

***S1 Video: Suppl_Groundthruth.mp4***

Original movie snippet from .*re_potemkin*, a copyleft crowdsourcing free/open source cinema project (https://re-potemkin.httpdot.net/).

***S2 Video: Suppl_Fig2E.mp4***

Reconstruction of the original movie snippet (Video 1).

***S3 Video: Suppl_Fig3B1.mp4***

Original movie with time extended in the middle.

***S4 Video: Suppl_Fig3B2.mp4***

As Supplemental Video 3 but with a new scene added post hoc.

***S5 Video: Suppl_Fig4B1.mp4***

Reconstruction of original movie (Video 1) with importance reduced in the last frames.

***S6 Video: Suppl_Fig4B2.mp4***

Reconstruction of original movie (Video 1) with importance reduced in the first frames.

***S7 Audiofile: Suppl_Fig6CFiltOrig.wav***

Filtered version (see Methods) of the original sound snippet song from the song *I’ll be your everything* by Texas Radio Fish (http://ccmixter.org/files/texasradiofish/63300, CC BY NC).

***S8 Audiofile:***

***Suppl_Fig6CReconstruction.wav***

Reconstruction of Supplemental Audio 1.

## Acknowledgments

The work was funded by the German Research Association (DFG) under Grant number LE2250/13-1. The author is indebted to Stefan Häusler for discussions and comments on the manuscript.

## Declarations

The author declares no competing interests.

## Repositories

The core recursive KSVR implementation can be found at https://github.com/cleibold/recsvr.

